# The Tip60/Ep400 chromatin remodeling complex impacts basic cellular functions in cranial neural crest-derived tissue during early orofacial development

**DOI:** 10.1101/2022.12.21.521531

**Authors:** Sebastian Gehlen-Breitbach, Theresa Schmid, Franziska Fröb, Gabriele Rodrian, Matthias Weider, Michael Wegner, Lina Gölz

## Abstract

The cranial neural crest plays a fundamental role in orofacial development and morphogenesis. Accordingly, mutations with impact on the cranial neural crest and its development lead to orofacial malformations such as cleft lip and palate. As a pluripotent and dynamic cell population, the cranial neural crest undergoes vast transcriptional and epigenomic alterations throughout the formation of facial structures pointing to an essential role of factors regulating chromatin state or transcription levels. Using CRISPR/Cas9-guided genome editing and conditional mutagenesis in the mouse, we here show that inactivation of Kat5 and Ep400 as the two essential enzymatic subunits of the Tip60/Ep400 chromatin remodeling complexes severely affects carbohydrate and amino acid metabolism in cranial neural crest cells. The resulting decrease in protein synthesis, proliferation and survival leads to a drastic reduction of cranial neural crest cells early in fetal development and a loss of most facial structures in the absence of either protein. Following heterozygous loss of Kat5 in neural crest cells palatogenesis was impaired. These findings point to a decisive role of the Tip60/Ep400 chromatin remodeling complex in facial morphogenesis and lead us to conclude that the orofacial clefting observed in patients with heterozygous KAT5 missense mutations is at least in part due to disturbances in the cranial neural crest.

## Introduction

Clefts of the lip and palate, jointly referred to as orofacial clefts, rank among the most prevalent human birth defects with a frequency of 1:700 ^1, 2^. They occur in syndromic or non-syndromic forms ^2, 3^. Both genetic alterations and environmental factors contribute to their emergence ^1, 4, 5^. Considering that proper orofacial development mainly depends on the ectodermal epithelium, the cranial neural crest-derived facial mesenchyme and the reciprocal interaction between both tissues, primary defects can be localized either to the ectodermal or the neural crest-derived tissue. It may affect any of a number of processes including precursor cell expansion, migration, differentiation, directional growth or fusion processes ^6–9^.

All of these processes are under tight control of a cell-specific gene regulatory network that consists of transcription factors, regulatory RNAs and chromatin modifying factors and is regulated by several signaling pathways in a tightly controlled temporospatial pattern. Among network components, chromatin modifying factors are particularly interesting, as they are both sensitive to genetic alterations and environmental factors. Previous studies have for instance pointed to important roles for components of the BAF and PBAF complexes and the Chd7 chromatin remodeler in orofacial development ^10–12^.

A recent study identified several de novo missense mutations in the chromodomain and the acetyl-CoA binding site of the histone acetyltransferase KAT5/TIP60 (henceforth referred to as KAT5) as cause for a syndromic form of orofacial clefting ^13^. The presence of other facial dysmorphisms points to a broader role of KAT5 in the neural crest-derived mesenchyme. Additionally, KAT5 has also been reported to be a quantitative trait locus and risk factor for non-syndromic forms of orofacial clefting because of the presence of single nucleotide polymorphisms in the vicinity of the gene ^14^.

Kat5 is a transcriptional co-activator that functions as part of the Tip60/Ep400 chromatin remodeling complex ^15, 16^. In this complex, Kat5 is responsible for acetylation of histones H2A and H4, whereas the Swi/Snf-type ATPase Ep400 catalyzes exchange of the acetylated H2A against the H2A.Z isoform that exhibits a preferential association with promoter regions and impacts gene expression. As other histone acetyltransferases, Kat5 may acetylate further proteins and have additional functions outside the Tip60/Ep400 complex ^17^. In agreement with their highly conserved function as chromatin modifiers, both Kat5 and Ep400 show a high degree of similarity between human and mouse with amino acid identities of 99% for Kat5 and 84% for Ep400.

Given the causal link with both syndromic and non-syndromic forms of orofacial clefts, we here decided to study the molecular function of Kat5 in the cranial neural crest and the facial mesenchyme. For conclusions about the Tip60/Ep400 complex, we extended our study to Ep400, the other central subunit of the complex.

## Results

### Kat5-dependent alterations of gene expression in a cranial neural crest cell line

To study the role of Kat5 in cells of the cranial neural crest, we first resorted to mouse O9-1 cells as a suitable and well characterized cellular model ^18^. In these cells, we used CRISPR/Cas9-dependent genome editing to inactivate the *Kat5* gene by targeting position 669 of the coding sequence in exon 8 (Fig. 1a). Position 669 is localized in the first common exon of *Kat5* immediately in front of the sequences coding for the MYST type HAT domain (Supl. Fig. 1a). According to Inference of CRISPR Edits (ICE) analysis, we succeeded in obtaining three independent clones that carried insertions or deletions in all four *Kat5* alleles of the tetraploid O9-1 cells (Suppl. Fig. 1b). These are predicted to lead to frameshifts and immediately ensuing premature translational termination (Suppl. Fig. 1b). Quantitative RT-PCR confirmed a drastic reduction of *Kat5* transcripts in all clones (Fig. 1b). In agreement, Kat5 protein was not detectable in any of the clones by immunocytochemical staining or by Western blotting of whole cell extracts (Fig. 1c,d; Suppl. Fig. 1c,d).

**Figure 1:**
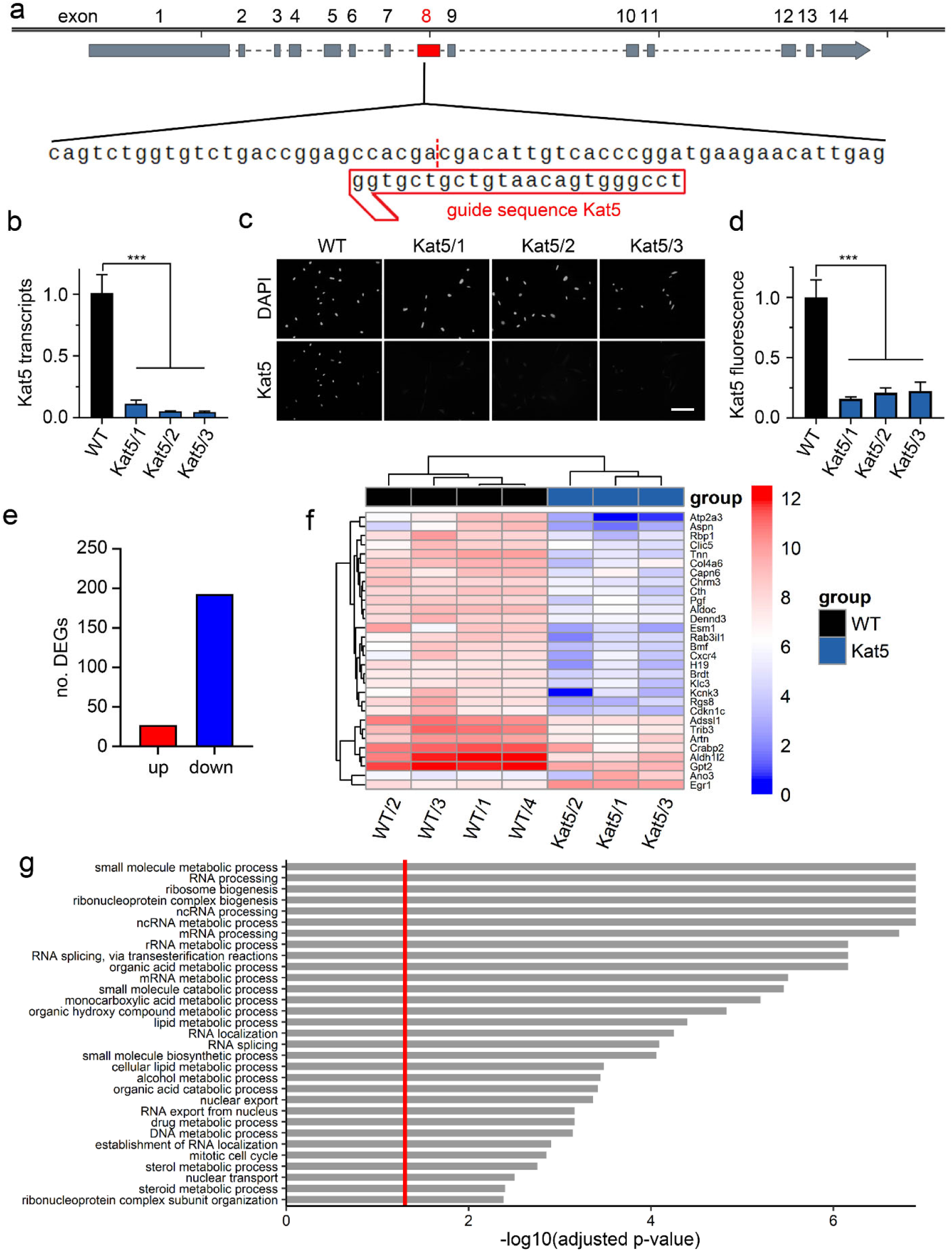
CRISPR/Cas9-dependent Kat5 inactivation in O9-1 cells. **(a)** Schematic representation of the *Kat5* gene with zoom-in on guide sequence and adjacent regions in the targeted exon 8 (red box). **(b)** *Kat5* transcript levels in wildtype O9-1 cells (WT) and gene-edited clones Kat5/1 – Kat5/3 as determined by RT-PCR (n=3). Levels in wildtype cells were set to 1 ± SEM. **(c,d)** Immunocytochemical detection of Kat5 protein in wildtype O9-1 cells (WT) and gene-edited clones (c), used for quantification (n = 3) of Kat5 levels (d). Nuclei were counterstained with DAPI. Scale bar: 150 µm. **(e-g)** Summary of the consequences of Kat5 inactivation in O9-1 cells as determined by RNA sequencing of gene-edited clones (n=3) and wildtype cells (n=4). Shown are the number of up-regulated (red) and down-regulated (blue) genes upon Kat5 inactivation (log2-fold ≥1, p-value ≤0.05) (e), a bi-clustering heatmap for visualization of the expression profile of the top 30 differentially expressed genes sorted and plotted by their (absolute) log2 transformed expression values in the samples (f), and a gene set enrichment analysis of preranked genes to identify cellular processes and structures affected by Kat5 inactivation (g). Statistical significance was determined by one-way ANOVA and Dunnett’s post test (***, P ≤0.001).

Using total RNA from the Kat5-deficient clones and O9-1 controls, we performed RNA sequencing studies. Kat5-deficient clones and controls clustered separately in PCA plots (Suppl. Fig. 1e). In total, we obtained 230 significantly deregulated genes (DEGs), of which 37 were upregulated and 193 were downregulated (absolute value of log2-fold ≥1, p-value ≤0.05) in the Kat5 knockout (ko) clones (Fig. 1e). Heatmaps of the 30 most strongly deregulated genes confirmed concordance among the samples in the respective groups (Fig. 1f). The proportion of up-versus downregulated genes is in accord with the proposed role of Kat5 as a general activator. Gene set enrichment analysis (GSEA) revealed that enriched sets were associated primarily with various aspects of the intermediate metabolism (e.g. small molecule biosynthesis/catabolism, organic hydroxyl compound, monocarboxylic acid and lipid metabolism), protein biosynthesis (e.g. ribosome biogenesis, RNA splicing, RNA nuclear export) and cellular proliferation (e.g. cell cycle, DNA replication, mitosis) (Fig. 1g). KEGG clustering of downregulated genes by process confirmed the strong alterations in gene sets associated to carbohydrate, lipid and amino acid metabolism and revealed links to several signaling pathways (Suppl. Fig. 1f).

### Ep400-dependent alterations of gene expression in a cranial neural crest cell line

To see how characteristic these alterations are for the Tip60/Ep400 complex in O9-1 cells, we next performed CRISPR/Cas9 dependent genome editing for the *Ep400* gene by targeting position 3201 of the coding sequence in exon 15 (Fig. 2a). Position 3201 is localized at the ATP binding site in the front part of the split ATPase domain of Ep400 (Suppl. Fig. 2a). We obtained one clone with insertions or deletions in all four *Ep400* alleles that led to frameshifts and premature translation termination (clone 1 in Suppl. Fig. 2b). A second clone contained a frameshift mutation in three alleles and a deletion of four amino acids in the remaining allele (clone 3 in Suppl. Fig. 2b). The third had a frameshift in one allele and a six amino acid deletion in the other three (clone 2 in Suppl. Fig. 2b). Amino acid deletions were all localized in the ATP binding site and should minimally lead to inactivation of Ep400. Quantitative RT-PCR again pointed to a drastic reduction of *Ep400* transcripts in all three clones (Fig. 2b). Additionally, immunocytochemical stainings failed to detect Ep400 in all clones arguing for an absence of the protein (Fig. 2c,d). Clustering of samples was again evident from the PCA plot (Suppl. Fig. 2c). In case of Ep400, we obtained a total number of 733 DEGs, 492 of them up- and 281 downregulated (absolute value of log2-fold ≥1, p-value ≤0.05) in the knockout clones (Fig. 2e). Heatmaps of the 30 most strongly deregulated genes again confirmed concordance among the samples in the respective groups (Fig. 2f). A comparable number of up- and downregulated genes has been observed in RNA sequencing analyses of other Ep400- containing and Ep400-deficient samples ^19–21^.

**Figure 2:**
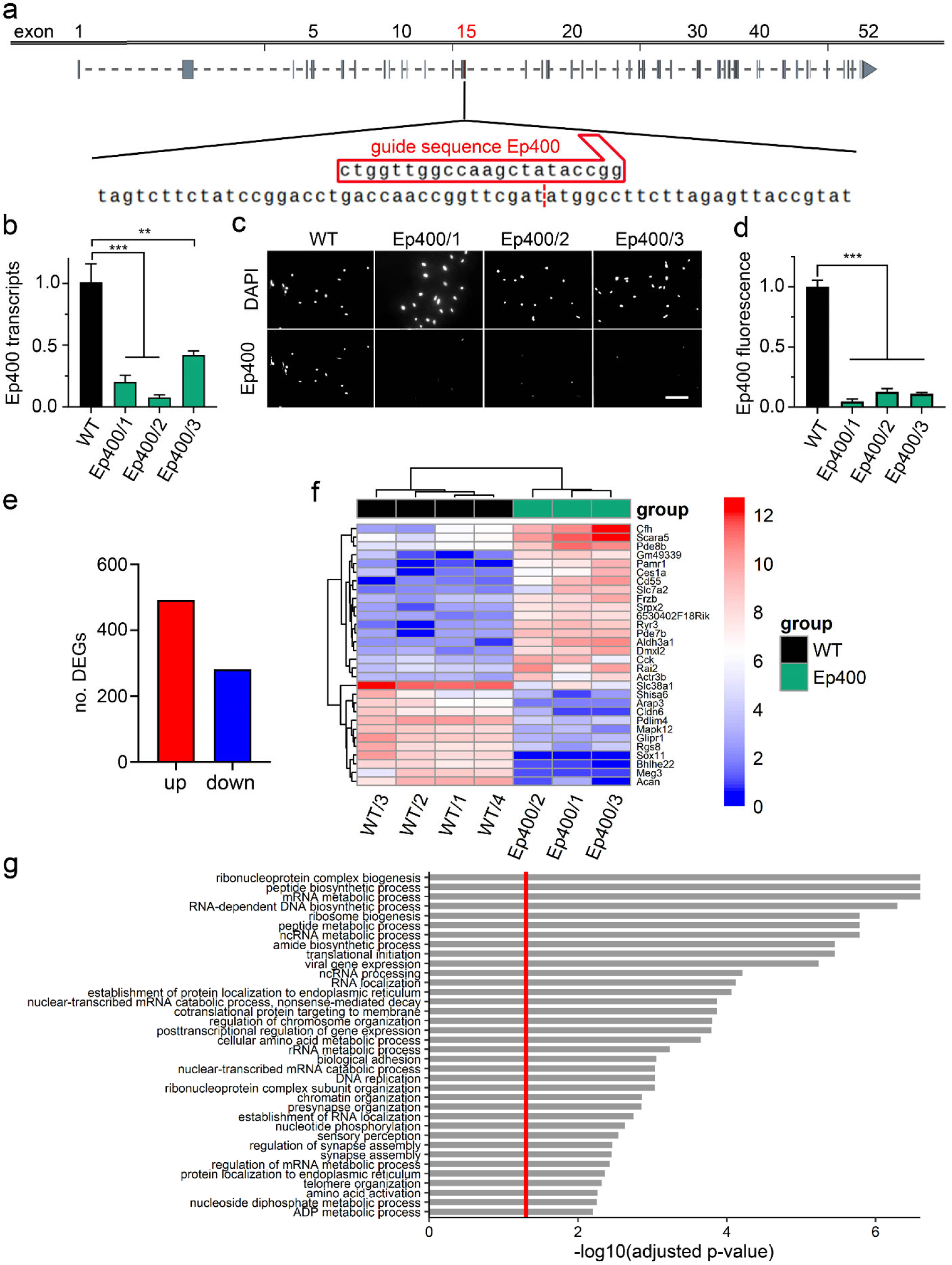
CRISPR/Cas9-dependent Ep400 inactivation in O9-1 cells. **(a)** Schematic representation of the *Ep400* gene with zoom-in on guide sequence and adjacent regions in the targeted exon 15 (red box). **(b)** *Ep400* transcript levels in wildtype O9-1 cells (WT) and gene-edited clones Ep4005/1 – Ep4005/3 as determined by RT-PCR (n=3). Levels in wildtype cells were set to 1 ± SEM. **(c,d)** Immunocytochemical detection of Ep400 protein in wildtype O9-1 cells (WT) and gene-edited clones (c), used for quantification (n = 3) of Ep400 levels (d). Nuclei were counterstained with DAPI. Scale bar: 150 µm. **(e-g)** Summary of the consequences of Ep400 inactivation in O9-1 cells as determined by RNA sequencing of gene-edited clones (n=3) and wildtype cells (n=4). Shown are the number of up-regulated (red) and down-regulated (blue) genes upon Ep400 inactivation (log2-fold ≥1, p-value ≤0.05) (e), a bi-clustering heatmap for visualization of the expression profile of the top 30 differentially expressed genes sorted and plotted by their (absolute) log2 transformed expression values in the samples (f), and a gene set enrichment analysis of preranked genes to identify cellular processes and structures affected by Ep400 inactivation (g). Statistical significance was determined by one-way ANOVA and Dunnett’s post test (**, P ≤0.01; ***, P ≤0.001).

Inactivation of the chromatin remodeling subunit of the Tip60/Ep400 complex predictably led to deregulation of genes associated with chromatin/chromosome organization in GSE analyses (Fig. 2g). Additionally, and similar to results on Kat5 ko clones, enriched gene sets in Ep400 ko clones were associated with small molecule metabolism (in particular amino acid metabolism) and aspects of protein biosynthesis (e.g. translation initiation, cotranslational ER targeting of proteins, peptide biosynthesis) according to GSEA. KEGG clustering by process confirmed that downregulation of gene expression in the absence of Ep400 affected amino acid metabolism and protein biosynthesis (via aminoacyl-tRNA biosynthesis) and again pointed to alterations in carbohydrate metabolism (glycolysis/gluconeogenesis) (Suppl. Fig. 2d).

Almost two thirds of the DEGs obtained for Kat5 were also deregulated in Ep400 ko clones. To compare DEG profiles in Kat5 and Ep400 ko clones in an unbiased manner, we used the rank–rank hypergeometric overlap (RRHO) algorithm ^22^. Our analysis revealed a strong and statistically significant overlap between gene-expression signatures that primarily concerned the downregulated genes in Kat5 and Ep400 ko clones (Fig. 3a). Application of the same algorithm to the terms from GSEA analysis shows a similarly high concordance for the terms with low p-value (Fig. 3b). KEGG pathway analysis argued that the common DEGs for Kat5 and Ep400 were involved in aspects of amino acid and carbohydrate metabolism as well as pluripotency/stem cell pathways (Fig. 3c). However, none of the DEGs listed in the latter category was a bona fide pluripotency gene. They rather represented genes associated more generally with pathways involved in precursor cell properties such as proliferation and increased protein synthesis.

**Figure 3:**
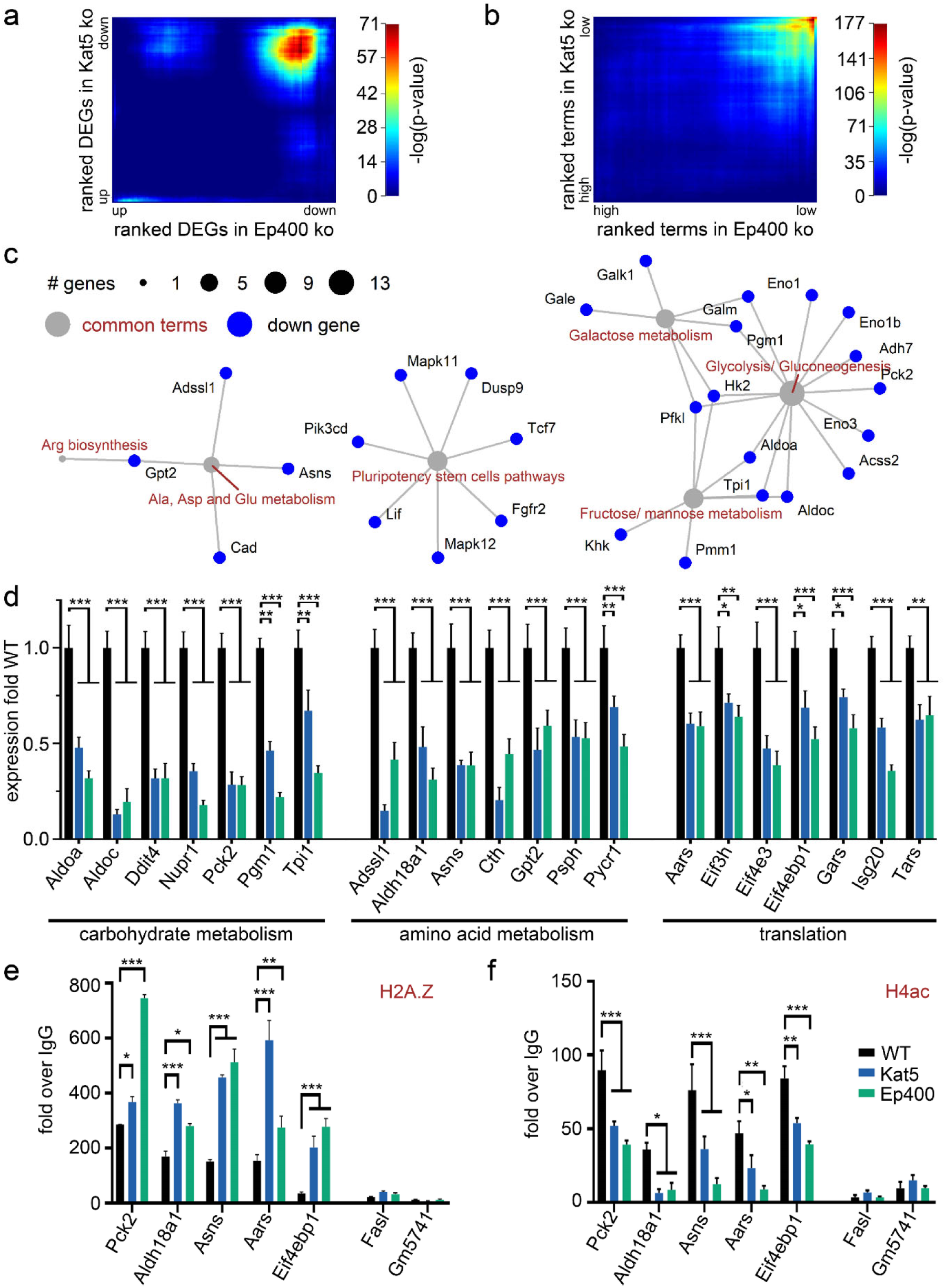
Common consequences of Kat5 and Ep400 inactivation on gene expression in O9-1 cells. **(a)** RRHO analysis for comparison of DEGs in Kat5 and Ep400 ko clones ranked by their degree of differential expression. The heatmap shows a strong and statistically significant overlap in the downregulated genes. **(b)** RRHO analysis of terms obtained by GSEA of preranked genes from Kat5 and Ep400 ko clones ranked by p-value reveals a strong overlap in terms with low p-value. **(c)** KEGG pathway analysis for the identification of the main cellular processes (grey dots, red terms) and the key components (blue dots, black terms) affected by both Kat5 and Ep400 inactivation. **(d)** Quantitative RT-PCR to validate expression changes of DEGs related to carbohydrate (left) and amino acid (middle) metabolism or protein biosynthesis (right) shared between Kat5 and Ep400 ko clones. Normalized transcript levels for each gene in wildtype O9-1 cells (WT) were set to 1, and levels in ko clones (n = 3) expressed relative to it. **(e,f)** Chromatin immunoprecipitation to determine the occupancy of H2A.Z (e) and acetylated H4 (H4ac, f) near the transcriptional start site of the *Pck2, Aldh18a1, Asns*, *Aars* and *Eif4ebp1* genes in wildtype O9-1 cells, Kat5 ko clone Kat5/3 and Ep400 clone Ep400/2. *Fasl1* and *Gm5741* genes served as controls. Amounts in precipitates were normalized to input and are presented as enrichment with histone-specific antibodies over IgG controls. Statistical significance was determined by 2-way ANOVA and Dunett’s post test (e,f) (*, P ≤0.05; **, P ≤0.01; ***, P ≤0.001).

A list of top downregulated genes categorized by their association with carbohydrate and amino acid metabolism as well as protein biosynthesis is presented in Suppl. Fig. 2e. For select DEGs, down-regulation in Kat5 and Ep400 ko clones was confirmed by quantitative RT-PCR (Fig. 3d). Interestingly, *Pck2, Aldh18a1, Asns, Aars*, and *Eif4ebp1* as representatives of genes associated with carbohydrate and amino acid metabolism or protein biosynthesis exhibited statistically significant increases in H2A.Z accumulation and statistically significant decreases in H4 acetylation near their transcriptional start sites in Kat5 and Ep400 ko clones according to chromatin immunoprecipitation analysis (Fig. 3e,f). In contrast, no comparable alterations were observed for the transcriptional start regions of the control genes *Fasl* and *Gm5741*. These data argue that at least some of the DEGs are direct and shared targets for Kat5 and Ep400 action.

### Impact of Kat5 and Ep400 on glycolysis, translation, proliferation and cell death in a cranial neural crest cell line

To evaluate the consequences of the observed expression changes on basic cellular functions in Kat5 ko and Ep400 ko clones, we analyzed several physiological parameters and compared them to the original O9-1 line. As a first step, we looked into the rate of ATP generation using the Seahorse XFe analyzer. In line with the RNA sequencing data and the inferred changes in gene expression, we measured lower ATP generation rates by glycolysis in the absence of Kat5 or Ep400 (Fig. 4a). This was consistent between all three clones with reductions ranging from 36% to 44% among Kat5 ko clones and from 34% to 56% among Ep400 ko clones (Suppl. Fig. 3a,b). In contrast, mitochondrial ATP generation by oxidative phosphorylation was not statistically different between Kat5 and Ep400 ko clones on the one and controls on the other side when averaged over all three clones (Suppl. Fig. 3c). At single clone level, one Kat5 ko clone and two Ep400 ko clones exhibited statistically significant reductions in mitochondrial ATP generation (Suppl. Fig. 3d,e). However, the respective levels of reduction were mild.

**Figure 4:**
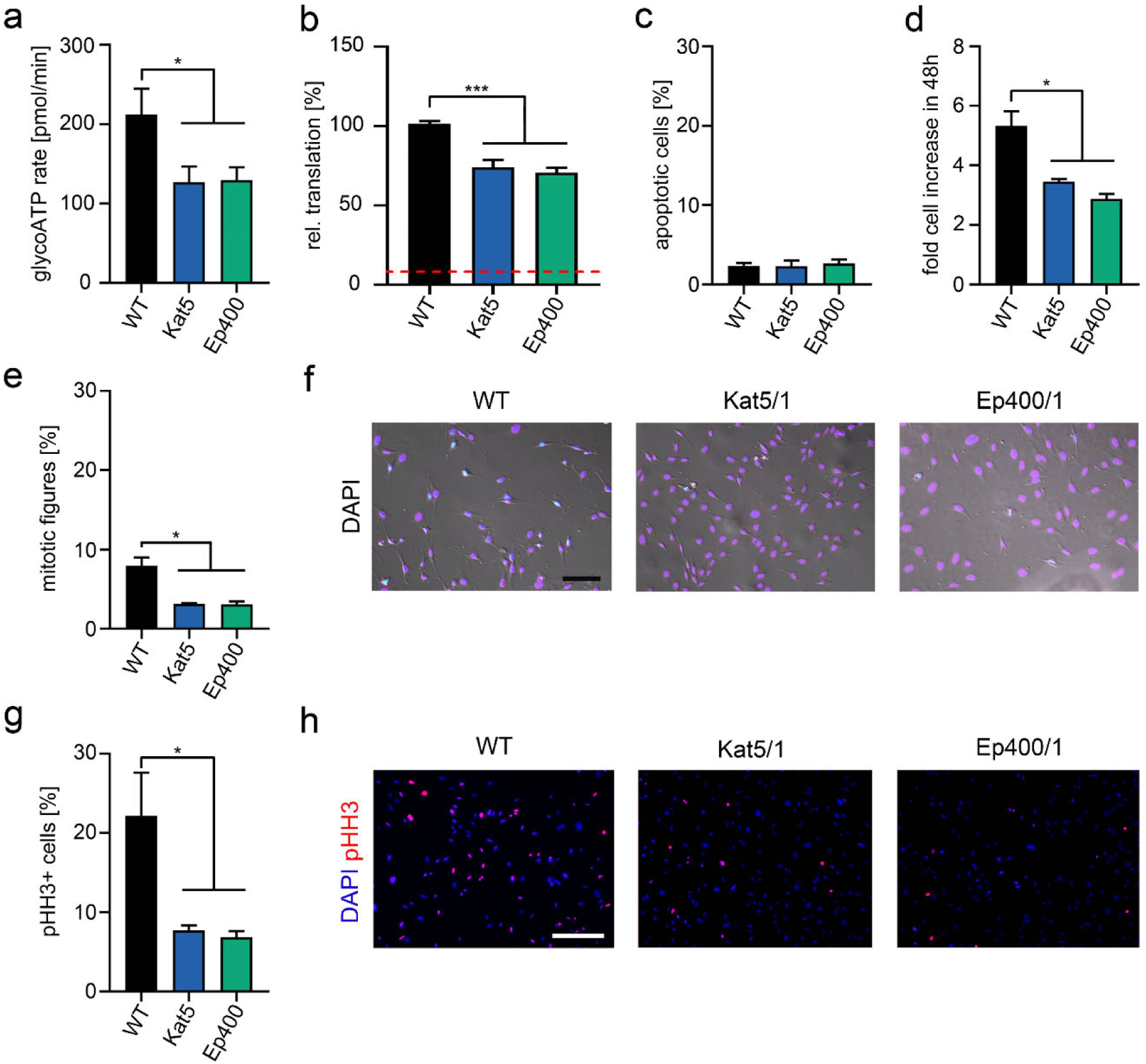
Altered basic functions in O9-1 cells upon Kat5 and Ep400 inactivation. **(a-d)** Comparison of O9-1 cells before (WT, black bars) and after inactivation of Kat5 (blue bars) or Ep400 (green bars) regarding rate of glycolytic ATP generation (determined in a Seahorse XFe analyzer as pmol per minute) (a), translation (determined as OPP incorporation in nascent transcripts with levels in WT cells set to 100%) (b), apoptosis (determined as percentage of cleaved caspase 3-positive cells among all cells) (c) and cell increase during 48 h (determined by crystal violet staining with staining at 0 h set to 1) (d). Red line in (b) corresponds to OPP incorporation in the presence of cycloheximide. Values were averaged over all three clones and are expressed as mean ± SEM. **(e-h)** Percentage of cells undergoing mitosis (e,f) or being positive for phospho-histone H3 (pHH3) (g,h) in 30% confluent cultures as additional parameters for proliferation. Quantifications (e,g) and representative phase contrast images with overlaid fluorescent DAPI signal (violet–blue to white; scale bar: 100 µm; f) or immunofluorescent images (pHH3 in red, DAPI in blue; scale bar: 200 µm; h) are shown. Statistical significance was determined by one-way ANOVA and Dunnett’s post test (*, P ≤0.05; ***, P ≤0.001).

To measure protein synthesis, incorporation of the puromycin analog OPP was used as a proxy for the rate of translation. Pretreatment of cells with the protein synthesis inhibitor cycloheximide was used to determine background levels (Fig. 4b, red dotted line) and analysis of control O9-1 cells defined the standard level of 100%. Intriguingly, both Kat5- and Ep400- deficient cells exhibited lower protein synthesis with 74 ± 4.8% of control levels averaged over all Kat5 ko clones and 71 ± 3.2% for Ep400 ko clones (Fig. 4b). All Kat5 ko clones and Ep400 ko clones consistently exhibited significantly decreased rates of translation (Suppl. Fig. 3f,g). Despite the decreased rate of protein synthesis, cell death was comparable between Kat5 ko and Ep400 ko clones on the one side and control O9-1 cells on the other both when averaged over all three clones (Fig. 4c) or analyzed in each single Kat5 ko and Ep400 ko clone (Suppl. Fig. 3h,i). However, proliferation was significantly lower in the absence of Kat5 or Ep400 when determined over a period of 48 h by crystal violet staining (Fig. 4d). Again, effects were consistent between ko clones with a 32-38% decrease for Kat5 ko clones and a 43-52% decrease for Ep400 ko clones (Suppl. Fig. 4a-c). This translated into a strongly reduced percentage of mitotic figures in cultures of Kat5- or Ep400-deficient O9-1 cells (Fig. 4e,f) with little variation among the ko clones for each gene (varying between 3.0% and 3.2% for Kat5 ko clones and between 2.4% and 3.7% for Ep400 ko clones as compared to 7.9% for the control O9-1 cells; Suppl. Fig. 4d,e). Qualitatively similar decreases were also detected in phosphohistone H3 stainings (Fig. 3g,h and Suppl. Fig. 4f,g). In summary, these results confirm that the changes in gene expression upon Kat5 or Ep400 deletion indeed result in decreased glycolysis and glycolytic ATP production, lowered protein biosynthesis and substantially reduced proliferation without detectable increases in cell death in vitro.

### Impact of Kat5 and Ep400 on the cranial neural crest and facial development in vivo

In the second set of experiments, we studied whether the Kat5- and Ep400-dependent changes are also relevant in cranial neural crest cells in vivo as they migrate into and form the mesenchyme of the first two pharyngeal arches that later give rise to most of the facial structures ^8, 23^. For this purpose, we crossed mice carrying a floxed *Kat5* or *Ep400* allele with the *Wnt1::Cre* transgene and a *Rosa26-stopflox-YFP* reporter to be able to target and identify neural-crest derived cells in the pharyngeal arches (Fig. 5a). Controls contained only the combination of *Wnt1::Cre* transgene and a *Rosa26-stopflox-YFP* reporter.

**Figure 5:**
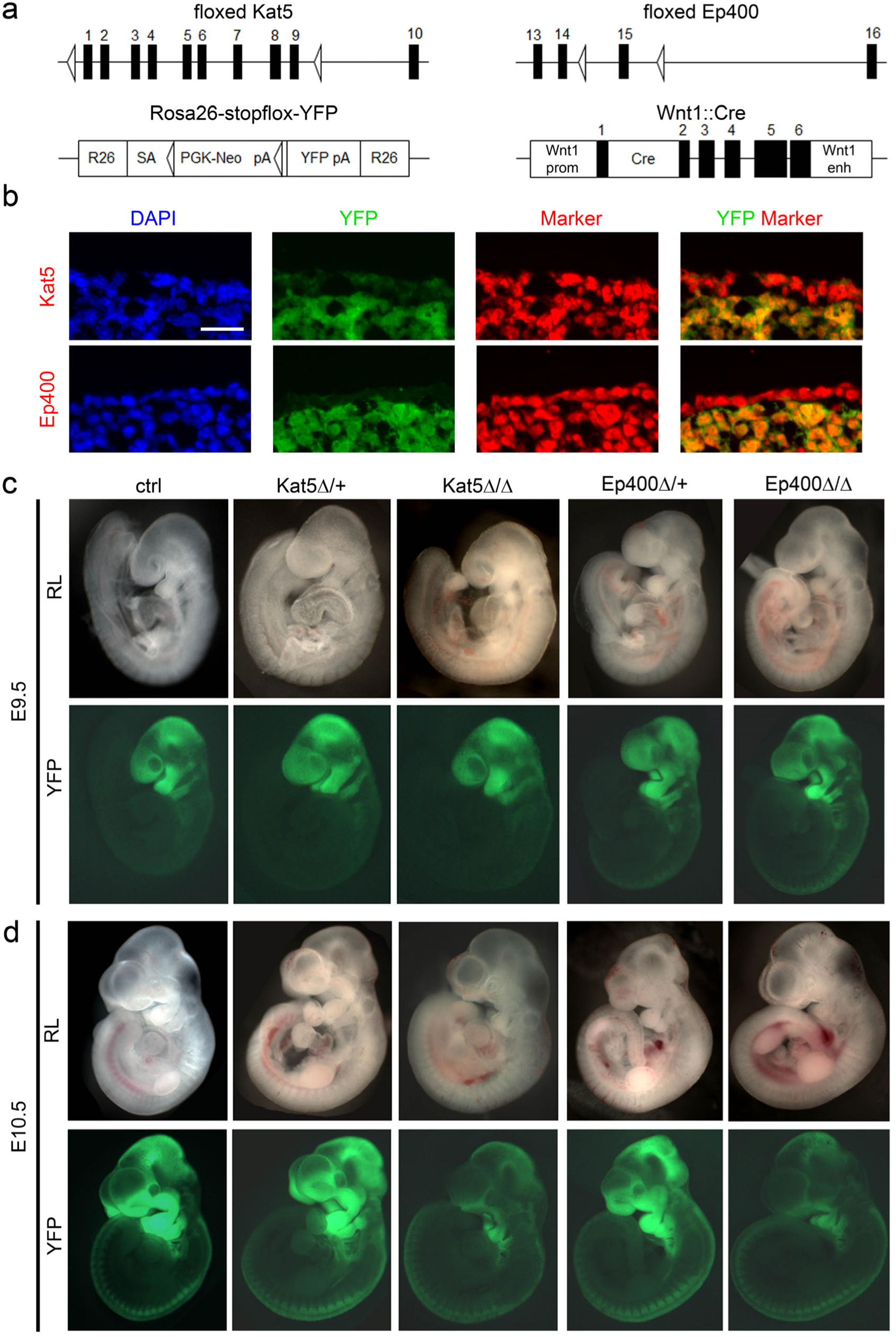
Developmental consequences of Kat5 and Ep400 inactivation in the cranial neural crest. **(a)** Schematic representation of the floxed alleles, the *Wnt1::Cre* used to inactivate *Kat5* and *Ep400* and the *Rosa26-stopflox-YFP* allele used to visualize successful inactivation in the cranial neural crest. Exons are depicted as black boxes, loxP sites as open triangles. pA, polyadenylation sequence; SA, adenovirus splice acceptor; PGK-Neo, pgk-neomycin; YFP, enhanced yellow fluorescent protein; Wnt1 prom, Wnt1 promoter; Cre, Cre recombinase; Wnt1 enh, Wnt1 enhancer. **(b)** Immunohistochemical staining of tissue from the first pharyngeal arch of control embryos at E10.5 with YFP (green) and Kat5 (red, upper row) or Ep400 (red, lower row). DAPI was used to counterstain nuclei. Scale bar: 25 µm. **(c-d)** Reflected-light microscopic (RL, upper rows) and YFP-autofluorescent (lower rows) images of whole mount control (ctrl) embryos and age-matched heterozygous as well as homozygous embryos with Kat5 (Kat5Δ/+, Kat5Δ/Δ) and Ep400 (Ep400Δ/+, Ep400Δ/Δ) deletions at E9.5 (c) and E10.5 (d).

Immunohistochemical staining of control tissue revealed that Kat5 and Ep400 proteins are similarly present and predominantly nuclear in the ectodermally-derived YFP-negative epidermis and the neural crest-derived YFP-positive mesenchyme of the pharyngeal arch (Fig. 5b). This argued for a comparable nuclear co-localization of Kat5 and Ep400 proteins in epidermal and neural crest-derived mesenchymal cells, but also underlined the necessity of conditional deletion to specifically study the role of Kat5 and Ep400 in the neural crest-derived cells.

Inspection of the whole mount embryos by reflected-light microscopy or autofluorescence revealed no major alterations or malformations at embryonic day (E) 9.5. This concerned the overall appearance, shape or size of the pharyngeal arches and the contribution of YFP-positive cranial neural crest cells in embryos lacking one or both copies of the *Kat5* (Kat5^Δ/+^ and Kat5^Δ/Δ^) or *Ep400* (Ep400^Δ/+^ and Ep400^Δ/Δ^) gene (Fig. 5c).

By E10.5, a reduced presence of neural crest cells in and contribution to the first two pharyngeal arches in Kat5^Δ/Δ^ and Ep400^Δ/Δ^ embryos was evident from the smaller size and altered morphology of the pharyngeal arches in whole mounts and the dramatically reduced YFP autofluorescence (Fig. 5d). In heterozygously deleted, i.e. Kat5^Δ/+^ and Ep400^Δ/+^ embryos, size and autofluorescent signal of pharyngeal arches remained relatively normal at E10.5.

In line with the impression from whole mount inspections, the total number of YFP-labeled neural crest-derived cells in the pharyngeal arch mesenchyme of Kat5^Δ/Δ^ and Ep400^Δ/Δ^ embryos was still fairly normal at E9.5 arguing against a major defect in early neural crest stem cell generation or migration in the embryos (Fig. 6a). By contrast, the number of cranial neural crest cells was strongly decreased in the first pharyngeal arch of Kat5^Δ/Δ^ and Ep400^Δ/Δ^ embryos at E10.5 (Fig. 6b,c). Cells in the pharyngeal arch of Kat5^Δ/Δ^ and Ep400^Δ/Δ^ embryos also displayed a significantly decreased expression of *Pck2, Tpi1, Aldh18a1, Gpt2, Aars* and *Eif3h* as genes involved in carbohydrate and amino acid metabolism or protein synthesis (Fig. 6d). This compares well with our previous observation in Kat5- and Ep400-deficient O9-1 cells (Fig. 3d). Proliferation rates were also substantially reduced in embryos with homozygous neural crest-specific deletion of *Kat5* or *Ep400* as determined by immunohistochemical staining for phosphohistone H3 (Fig. 6e,f; for representative stainings see Suppl. Fig. 5a-c). Intriguingly, a reduction in phosphohistone H3-positive cells was already visible at E9.5 (Fig. 6e; for representative stainings see Suppl. Fig. 5b). The reduced proliferation rates in Kat5^Δ/Δ^ and Ep400^Δ/Δ^ embryos were also confirmed in BrdU incorporation studies at E10.5. One hour after treatment, only 7-18% of neural crest cells had incorporated BrdU as compared to 44% in controls (Fig. 6g; Suppl. Fig. 5d).

**Figure 6:**
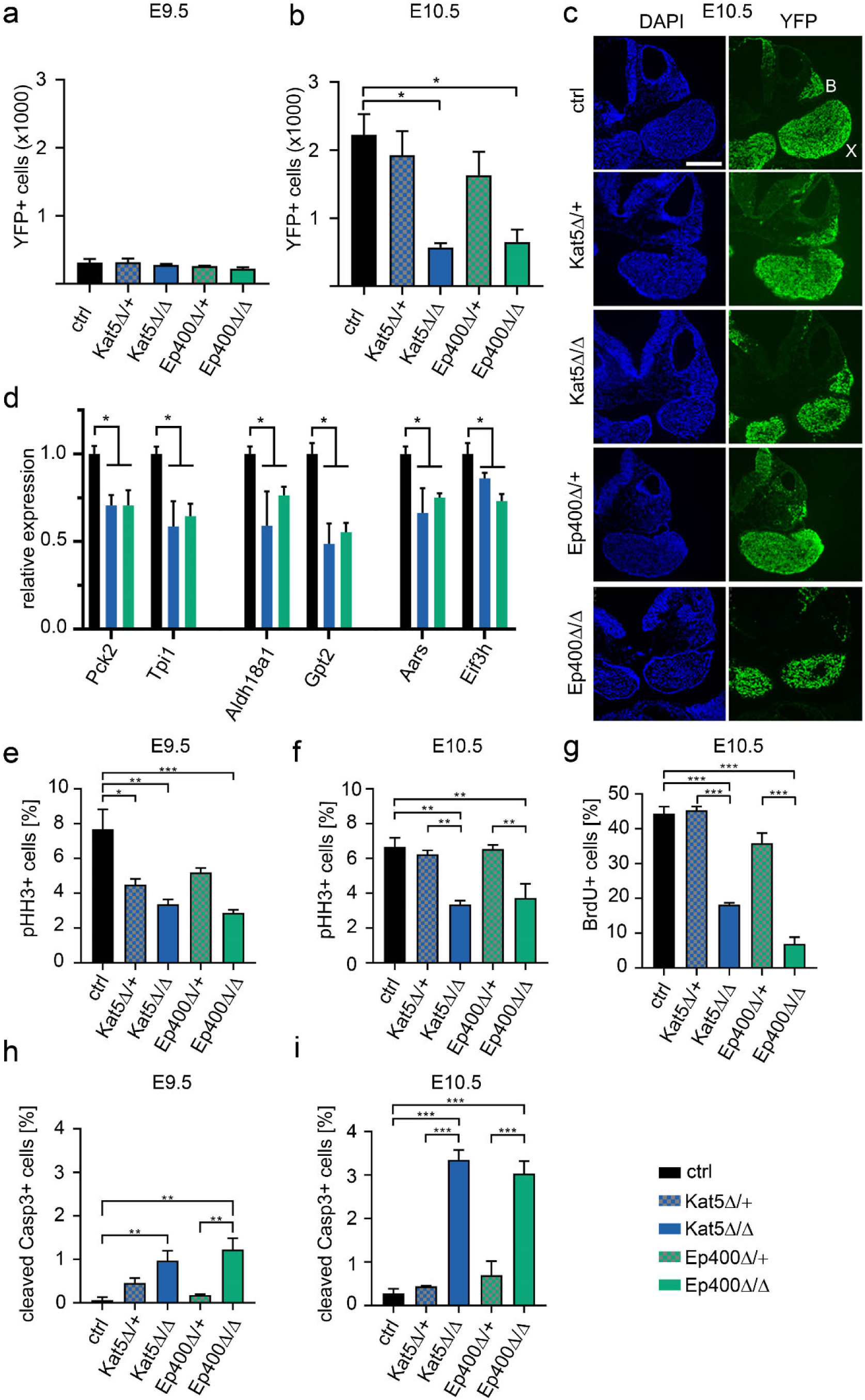
Consequences of Kat5 and Ep400 inactivation on gene expression, proliferation and survival of the developing cranial neural crest. **(a,b)** Overall numbers of YFP-labelled cells in the first pharyngeal arches of control (ctrl) embryos and age-matched heterozygous as well as homozygous embryos (n=3 for each genotype) with Kat5 (Kat5Δ/+, checkered blue; Kat5Δ/Δ, blue) and Ep400 (Ep400Δ/+, checkered green; Ep400Δ/Δ, green) deletions at E9.5 (a) and E10.5 (b). Absolute numbers ± SEM correspond to cells per first pharyngeal arch. **(c)** DAPI stain (left) and YFP-autofluorescence (right) of transverse sections from control, heterozygous and homozygous embryos with Kat5 or Ep400 deletions at E10.5 showing maxillary (X) and mandibular (B) branch of the first pharyngeal arch. Scale bar: 300 µm. **(d)** Quantitative RT-PCR to determine transcript levels of *Pck2, Tpi1, Aldh18a1, Gpt2, Aars* and *Eif3h* in pharyngeal arches of ctrl, Kat5Δ/Δ and Ep400Δ/Δ embryos at E10.5. Normalized transcript levels for each gene in ctrl embryos were set to 1, and levels in Kat5Δ/Δ and Ep400Δ/Δ embryos (n = 3) were expressed relative to it. **(e-g)** Percentages of phosphohistone H3-(pHH3-) positive (e,f) and BrdU-labelled (g) neural crest cells in the first pharyngeal arches of control, hetero- and homozygous embryos (n=3 for each genotype). Percentages are given as mean value ± SEM. **(h,i)** Rate of cleaved caspase 3-positive apoptotic cells in the various genotypes at E9.5 (h) and E10.5 (i). Statistical significance was determined by multiple t-test (d) or one-way ANOVA and Sidak’s post test (e-i) (*, P ≤0.05; **, P ≤0.01; ***, P ≤0.001).

It is noteworthy that even Kat5^Δ/+^ and Ep400^Δ/+^ embryos exhibited reduced numbers of phosphohistone H3-positive cells at E9.5 (Fig. 6e; for representative stainings see Suppl. Fig. 5b). This correlated with a lower number of neural crest-derived cells in the pharyngeal arch mesenchyme at E10.5, although the reduction in cell number was not statistically significant. By E10.5, the reduction of phosphohistone H3-positive cells in Kat5^Δ/+^ and Ep400^Δ/+^ embryos had disappeared (Fig. 6f; for representative stainings see Suppl. Fig. 5c). Nevertheless, this argues that even heterozygous loss of Kat5 or Ep400 may have mild effects on neural crest proliferation.

In contrast to O9-1 cells in vitro, neural crest cells with homozygously deleted *Kat5* or *Ep400* additionally exhibited increased rates of apoptosis as determined by immunohistochemical stainings for cleaved caspase 3 at both E9.5 and E10.5 (Fig. 6h,i; Suppl. Fig. 6a,b). Heterozygous deletion in Kat5^Δ/+^ and Ep400^Δ/+^ embryos also led to slightly increased cell death rates that did not reach statistical significance.

As a consequence of reduced proliferation and increased apoptosis, neural crest cell numbers were decreased by 70 – 75% in the first pharyngeal arch of Kat5^Δ/Δ^ and Ep400^Δ/Δ^ embryos at E10.5. In Kat5^Δ/+^ and Ep400^Δ/+^ embryos, neural crest cell numbers in the pharyngeal arches were only slightly decreased.

When analyzed shortly before birth at E18.5, the early cranial neural crest defect had translated into a near absence of facial structures in embryos with homozygous *Kat5* deletion (Fig. 7a). Similar gross facial malformations were also observed in age-matched embryos with homozygous *Ep400* deletion (Suppl. Fig. 6c), whereas neither mice with heterozygous *Kat5* deletion nor mice with heterozygous *Ep400* deletion exhibited any obvious facial malformations upon visual inspection (data not shown).

**Figure 7:**
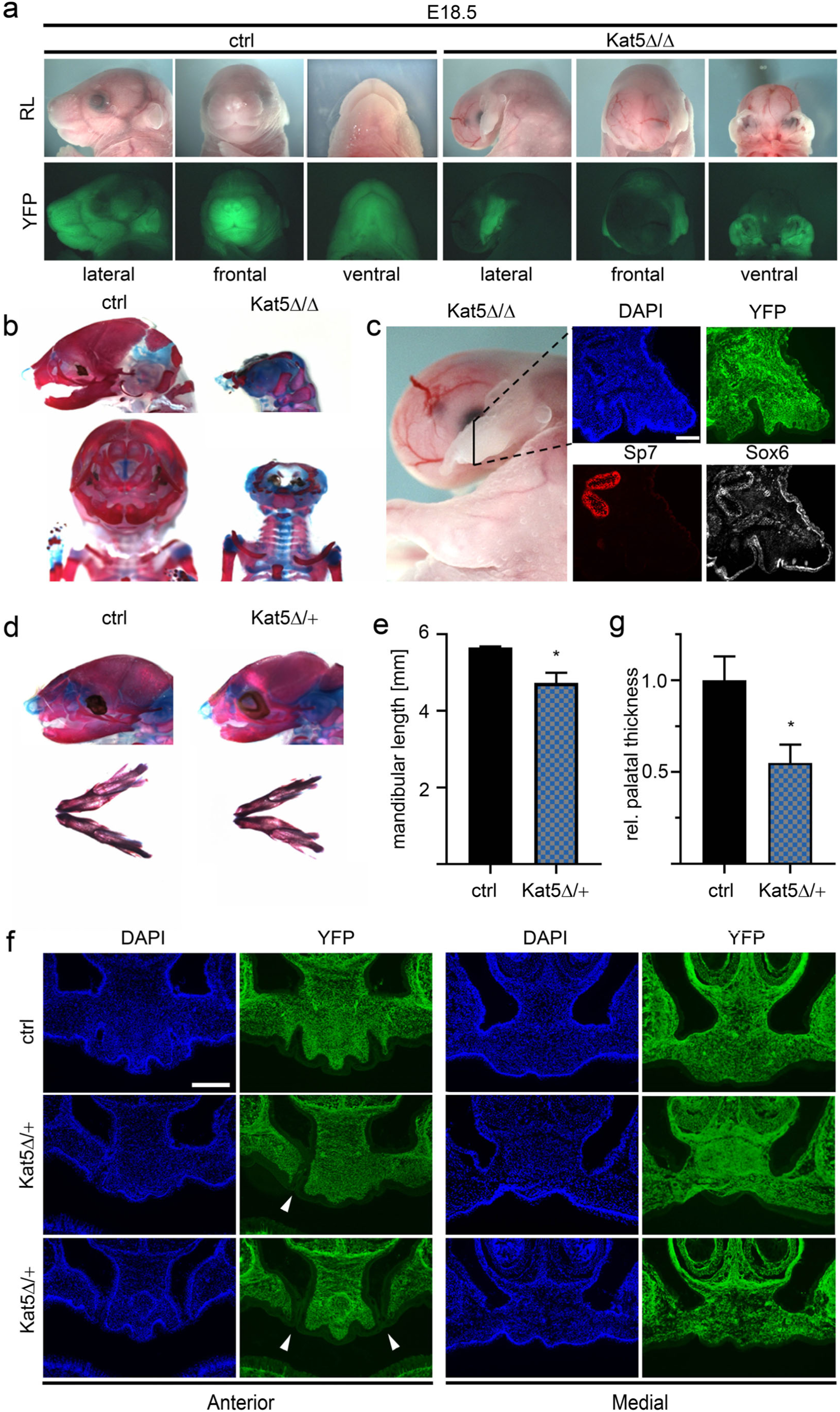
Perinatal phenotype of mice with neural crest-specific Kat5 inactivation. **(a)** Reflected-light microscopic (RL, upper row) and YFP-autofluorescent (lower row) images of heads from control (ctrl) embryos and age-matched homozygous embryos with Kat5 deletion (Kat5Δ/Δ) at E18.5. **(b)** Alcian blue/alizarin red stainings of cartilaginous and osseous head structures in control and Kat5Δ/Δ embryos at E18.5. **(c)** Immunohistochemical staining of the remaining facial structures in Kat5Δ/Δ embryos at E18.5 with antibodies directed against YFP (green), Sp7 (red) and Sox6 (white). Nuclei were counterstained with DAPI (blue). For overview of the structure and orientation of the plane of sectioning, see left panel. **(d)** Alcian blue/alizarin red stainings of head (top) and mandibles (bottom) from control and Kat5Δ/+ embryos at E18.5. **(e)** Determination of mandibular length in control and Kat5Δ/+ embryos at E18.5 (n = 3). **(f)** Visualization of the anterior (left) and middle (right) part of the secondary palate of control and Kat5Δ/+ embryos at E18.5 by DAPI and YFP staining of coronal sections. Arrowheads point to clefts in the secondary palate. Scale bars: 200 µm (c,f). **(g)** Quantification of secondary palatal thickness (middle region) in control and Kat5Δ/+ embryos at E18.5 (n = 3). Statistical significance was determined by unpaired t-test (*, P ≤0.05).

A combined alcian blue/alizarin red staining of Kat5^Δ/Δ^ embryos revealed a drastic reduction of cartilage and bone in the facial region (Fig. 7b). However, some osseous and cartilaginous structures remained. Due to the strong malformations, it was impossible to assign the remnants to any particular facial bone or cartilage. However, immunohistochemistry verified that Sp7- positive bone-forming cells and Sox6-positive chondrocytes were present (Fig. 7c). Each group of cells labelled additionally with the YFP lineage marker, attesting to their neural crest origin (Fig. 7c). We therefore conclude that neither osteogenic nor chondrogenic differentiation is completely blocked.

A closer look at the facial skeleton of Kat5^Δ/+^ embryos revealed no missing structures. However, the mandible was significantly shortened (Fig. 7d,e). Histochemical analyses with DAPI and YFP revealed the presence of unilateral or bilateral clefts in the anterior most regions of the secondary palate in a subset of the embryos (Fig. 7f, left). From the middle to the most posterior region, the secondary palate was fused, but appeared substantially thinner than its counterpart from age-matched controls (Fig. 7f, right and Fig. 7g). These findings argue that a heterozyogous loss of *Kat5* in the cranial neural crest is sufficient to induce mild orofacial clefting.

## Discussion

In this manuscript, we have shown the relevance of Kat5 and Ep400 as the two central components of the Tip60/Ep400 chromatin remodeling complex during early development of the neural crest-derived facial mesenchyme in vitro using the O9-1 neural crest-like cell line and in vivo using conditional mouse models. In particular, we have shown that both proteins have a strong impact on neural crest cell metabolism, especially on carbohydrate usage and amino acids availability.

The influence of Kat5 and Ep400 on metabolic processes was detected in RNA sequencing studies on genome edited O9-1 cells lacking either Kat5 or Ep400. The magnitude of deregulated genes and the ratio of upregulated versus downregulated genes were roughly comparable with other studies on the two factors in different cellular systems ^19–21, 24^. There was furthermore a substantial overlap in DEGs between the RNA sequencing results for the two factors. Additionally, bioinformatic analyses pointed to the fact that both Kat5 and Ep400 influence the same cellular processes and are thus functionally linked in O9-1 neural crest cells. A closer study of select DEGs in O9-1 cells furthermore revealed that they exhibited alterations in H2A.Z occupancy and H4 acetylation around their transcriptional start sites. This argues that at least some DEGs are direct and shared targets of Kat5 and Ep400, and thus likely under the influence of the Tip60/Ep400 complex. It also implies that loss of the histone modifying and chromatin remodeling activities of the Tip60/Ep400 complex underlies the observed phenotypes. Some DEGs that were identified in O9-1 cells were similarly altered in their expression upon Kat5 and Ep400 inactivation in cranial neural crest cells in vivo. Therefore, it seems plausible to assume that the results from O9-1 cells are at least in part also relevant in vivo.

A functional linkage between Kat5 and Ep400 is also supported by the very similar behavior of Kat5- and Ep400-deficient O9-1 cells in our functional assays and by the very similar phenotypic alterations in the neural crest-specific Kat5 and Ep400 mouse mutants.

In line with the alterations in carbohydrate and amino acid metabolism, glycolytic ATP generation and protein biosynthesis are strongly impaired. This almost certainly contributes to the substantially decreased proliferation rate of Kat5- or Ep400-deficient cells. Proliferation has been reported to be reduced after deletion of Kat5 or Ep400 in some cell types (e.g. fibroblasts), to be increased in others (e.g. cardiomyocytes) or remained unaltered (e.g. Schwann cells, oligodendroglia) ^20, 25–28^. This argues that the impact of the Tip60/Ep400 complex on proliferation is highly cell type-specific.

Additionally, we detected a substantially higher cell death rate in cranial neural crest cells in vivo, but not in vitro. We assume that the cells are better able to cope with metabolic impediments in tissue culture because of the excess of nutrients provided in the medium and the possibility to switch to higher mitochondrial ATP generation. In vivo, however, migrating neural crest cells constitutively rely for energy production on aerobic glycolysis ^29^. Without the ability to compensate for the energy shortage, cells may be more susceptible to cell death. As a result of the decreased proliferation and increased cell death, cranial neural crest cells fail to form sufficient amounts of mesenchyme in the pharyngeal arches that later constitute the facial mesenchyme and are required for proper formation of primary and secondary palate and final fusion processes. The resulting dramatic facial malformations and orofacial clefting in mice with homozygous neural crest-specific deletion of Kat5 or Ep400 around the time of birth are an impressive confirmation of the essential role of both gene products in facial development, including palatogenesis.

Similarly dramatic disruptions of facial development have been observed in other neural-crest specific mouse mutants including mice with loss of β-catenin or ephrin-B1/2 ^30–32^. Whether there is any direct link between the roles of Kat5 and Ep400 on the one side and the affected gene products on the other is currently unknown.

Intriguingly, we also see mild alterations in the number and proliferation of cranial neural crest cells in mice with heterozygous cell-specific deletion of *Kat5* and *Ep400*. At least in mice with heterozygous neural crest-specific deletion of *Kat5* this seems to translate into mild clefts in the secondary palate. Currently we do not know whether similar clefts occur in mice with heterozygous neural crest-specific deletion of *Ep400,* but would expect this to be the case.

On the basis of the findings presented in this study, it is reasonable to assume that the early defects in the survival and expansion of the cranial neural crest are also at least in part causative for the syndromic orofacial clefts observed in patients with heterozygous Kat5 mutations and may also explain the role of Kat5 as a quantitative trait locus in non-syndromic forms ^13, 14^.

Orofacial clefts in patients with heterozygous Kat5 mutations appear to be more severe than those observed in mice. The most parsimonious explanation for the different severity are quantitative or qualitative species-specific differences in facial development. Already due to size differences, many more neural crest cells will be needed in humans than in mice for generating sufficient amounts of facial mesenchyme. Alternatively, it cannot be excluded that there is an additional function of Kat5 on epidermal development that needs to be impaired for phenotypic alterations to become visible.

## Conclusion

Our data show that Kat5 and Ep400 as two components of the Tip60/Ep400 chromatin remodeling complex play an important role in the metabolic reprogramming of neural crest cells and influence survival and proliferation. As a consequence, *Kat5* and *Ep400* mutations impair development of the neural crest-derived facial structures. In case of heterozygous neural crest-specific *Kat5* mutations, this leads to orofacial clefts. As the activities of Kat5 and Ep400 are themselves impacted by key metabolites such as acetyl-CoA and ATP, they may play a central role in sensing the metabolic status and in metabolically reprogramming the cells by alterations in chromatin changes. This needs to be investigated in future studies, but may eventually be instrumental in manipulating the cranial neural crest in a therapeutically relevant manner.

## Materials & Methods

### Cell Culture

The murine neural crest cell line O9-1 was obtained from Sigma Aldrich and cultured as published ^33^. Briefly, O9-1 cells were maintained on Matrigel-coated dishes in preconditioned O9-1 medium. O9-1 medium consisted of DMEM (Gibco) supplemented with 15% FCS, 0.1 mM nonessential amino acids, 0.55 µM β-mercaptoethanol, 100 U/ml penicillin and 100 U/ml streptomycin. For preconditioning, O9-1 medium was incubated with mitomycin C (Adooq Bioscience) inactivated STO mouse fibroblasts (ATCC) for 24 h, sterile-filtered (Steritop, 0.22 µm) and supplemented with 1000 units/ml LIF, and 25 ng/ml FGF2. The O9-1 cells were passaged every three days and reseeded at a density of 1:40.

### Generation of CRISPR Knockout Lines

*Kat5* and *Ep400* deficient O9-1 cell lines were obtained by CRISPR/Cas9 mediated genome editing. Guide sequences (see Fig. 1a, Fig. 2a) were determined using the CRISPOR online server (Version 4.99) ^34^, shortened to 19 bp length and flanked with BbsI overhangs. Corresponding oligonucleotides (Suppl. Table 1) were inserted into BbsI-digested pX330-U6- Chimeric_BB-CBh-hSpCas9 (obtained from Addgene, plasmid #42230) ^35^ to generate pX330- Ep400-Ex15 and px330-Kat5-Ex8. These plasmids were co-transfected with pEGFP-N1 (Clontech) into O9-1 using Lipofectamine 2000 (Thermo Fisher). After 72 h, FACSorted EGFP-positive cells were individually seeded into 96-well plates, expanded to clones and checked for Ep400 or Kat5 expression by RT-PCR, immunocytochemistry and Western blotting. Additionally, genomic DNA around the targeted site was amplified by PCR (primer sequences see Suppl. Tables 1,2) and sequenced to determine a knockout score using Synthego ICE Analysis 2019, v3.0. Only clones with a score > 90% were selected.

### Western blotting

Total protein lysates were generated from 80% confluent O9-1 cultures by lysis in PE-buffer (5M urea, 10% glycerol, 0.1% SDS, 0.01 M Tris pH 6.8) with cOmplete Mini protease inhibitor (Sigma-Aldrich) and sonication. Protein concentrations were determined using the BioRad DC assay. After denaturation, 20 µg of protein were electrophoresed per lane alongside a molecular weight marker on SDS-polyacrylamide gels before blotting onto nitrocellulose membranes. Membranes were washed and blocked in 1x TBST with 5% bovine serum albumin before consecutive incubation with primary antibody (see Suppl. Table 3) and horseradish peroxidase-coupled protein A (Cymed) and detection using Clarity Western ECL Substrate (BioRad).

### Real Time PCR (RT-PCR)

Total RNA was extracted from O9-1 knockout clones at passage 2 after FACSorting and wildtype cells or from pharyngeal arches dissected from embryos at E10.5 using the RNEasy micro kit according to the manufacturer’s protocol (Quiagen). To obtain cDNA and validate expression levels, RNA was reverse-transcribed as described ^36^. PCR reactions were run on a CFX96 Touch Real-Time PCR Detection System (BioRad) with the resulting cDNAs in Power SYBR Green Mastermix (Thermo Fisher) and 125 nmol transcript-specific primers (see Suppl. Table 2). Ct values were determined using Bio-Rad CFX manager software and used to calculate fold changes by the ΔΔCt method.

### Chromatin Immunoprecipitation

O9-1 cells and knockout clones were treated at room temperature with 1% paraformaldehyde, lysed and used to prepare chromatin. Cross-linked chromatin was subsequently sheared with a Bioruptor (Diagenode) to fragments of 150-400 bp, precleared and incubated with 2µg rabbit anti-H2A.Z antibodies, rabbit anti-acetylated H4 antibodies or rabbit IgG (Suppl. Table 3). Following precipitation with BSA-blocked protein A sepharose beads, the precipitated chromatin underwent crosslink-reversal at 68°C in the presence of proteinase K and RNAse. From the reaction, DNA was purified with the NucleoSpin Gel and PCR cleanup Kit (Macherey-Nagel) and used for quantitative PCR to detect gene-specific regions near the respective transcriptional start (Suppl. Table 1). All samples were processed as technical triplicates. The ΔΔCt method was used to calculate the enrichment obtained in the precipitate with anti-histone antibodies over IgG.

### Immunocytochemistry

O9-1 cells were cultivated on Matrigel (Corning) coated and HCl treated glass coverslips before fixation in PBS containing 4% paraformaldehyde, permeabilization with 0.5% Triton X-100, blocking in 10% fetal calf serum, consecutive incubation with primary and secondary antibodies (see Suppl. Table 3), counterstaining of the nucleus with DAPI and mounting onto slides using Mowiol 4-88 (Carl Roth).

### RNA Sequencing and bioinformatic analysis

RNA of 4 biologically independent O9-1 wildtype samples as well as 3 different Kat5 and Ep400 knockout clones was used for library generation, poly-A enrichment and RNA sequencing on the NovaSeq 6000 platform with a read length of 2 x 100 bp and an output of ∼30 M clusters per sample (CeGaT, Tübingen). The generated sequence data were mapped onto the *Mus musculus* reference genome mm10 and normalized read counts were generated.

Bioinformatic analysis was performed using R version 4.2.1. The Bioconductor package *DESeq2* (version 1.36.0) ^37^ was applied for differential expression analysis with default parameters. To assess the similarity of the gene deregulation profile of Ep400 and Kat5 ko clones, a hypergeometric overlap approach was conducted on ranked logarithmic fold changes using R package RRHO (version 1.36.0) ^22^. The R package fgsea (version 1.22.0) ^38^ was used for gene set enrichment analysis using the complete gene lists preranked by logarithmic fold change with a false discovery rate (FDR) of less than 0.05. The requirements for deregulation were an absolute value of the logarithmic fold change of at least 1 and a group counts’ mean larger than 100. ReViGo (http://revigo.irb.hr/) ^39^ was used to summarize and visualize the results. R package RRHO (version 1.36.0) was applied to rank and compare the enriched GSEA gene sets, R package pathfindR (version 1.6.4) ^40^ to identify enriched Kyoto Encyclopedia of Genes and Genomes (KEGG) pathways in the set of deregulated genes.

### Crystal Violet Staining

O9-1 cells were seeded in triplicates at a density of 10,000 cells/cm^2^ on Matrigel coated dishes. One set of cells underwent fixation 2 h after seeding while a second identically seeded plate was incubated for 48 h at 37°C, 5% CO2. Fixation was performed with 70% ethanol for 30 min at 4°C. After fixation, the cells were stained for 30 min at room temperature with crystal violet staining solution, prepared by diluting 0.5% crystal violet (Carl Roth) in 20% methanol, and consecutively washed 5 times with ddH_2_O. Plates were dried overnight and the crystal violet was extracted with 0.1 M sodium citrate in 50% ethanol for 30 min at room temperature. Absorbance was measured at a wavelength of 550 nm and proliferation rates were calculated by dividing the absorbance after 48 h by the absorbance after 2 h for corresponding wells.

### Apoptosis Assay

Apoptosis of O9-1 cells was determined using propidium iodide (PI) (Fluka) staining followed by flow cytometric analysis ^41^. Briefly, O9-1 cells were cultured to 70% confluency, detached, fixed with 70% ice cold ethanol for 2 h, washed two times with PBS and incubated with an extraction buffer consisting of PBS with 0.1 M Na_2_HPO_4_ and 0.05% Triton X-100 for 5 min at room temperature. After centrifugation at 400 g for 5 min, cells were resuspended in PBS with 20 µg/ml PI and 0.2 mg/mL RNAse A (Macherey-Nagel) and incubated at RT for 30 min before analysis on a FACSCalibur (Beckton Dickinson). The rate of apoptosis was determined by calculating the fraction of cells with a sub-G1 DNA content using the Flowing Software 2.51 (Turku Bioscience Centre).

### Seahorse ATP-Rate Assay

Metabolic profiling of O9-1 cells was performed using a Seahorse XFe Analyzer (Agilent). O9-1 cells were seeded in preconditioned medium into Seahorse XFe 96 well plates at a density of 20,000 cells/well and incubated overnight. Before the measurement, the cells were washed twice and incubated in fresh XF DMEM pH7.4 (Agilent) supplemented with 10 mM glucose (Carl Roth), 1 mM sodium pyruvate (Gibco) and 2 mM glutamic acid (Gibco) for 45 min at 37°C without CO_2_. The cartridge was hydrated with XF Calibrant solution (Agilent) for 16 h, loaded with 75 µM oligomycin and 25 µM rotenone + antimycin A before measurement according to Agilent’s ATP-Rate assay protocol. After the measurement, the cells were washed with PBS, treated with 70% ethanol for 30 min and stained with crystal violet. The obtained cell density values were used for data normalization. Total ATP generation as well as mitochondrial and glycolytic contributions were determined using the Wave Desktop Application (Agilent) for the conducted assay.

### Protein Synthesis Assay

Protein synthesis was determined by measuring the incorporation of O-propargyl-puromycin (OPP) into newly synthesized proteins ^42^ using the Click-it Plus OPP Alexa Fluor 647 Proteinsynthesis-Assay-Kit according to the manufacturer’s manual (Thermo Fisher). Briefly, O9-1 cells were cultured in 96-well plates until 50% confluency and a pulse of OPP was applied for 30 min. Untreated cells served as negative control, cells pre-incubated with 50 µg/ml cycloheximide for 15 min defined baseline levels. After the incubation time, cells underwent fixation in 4% paraformaldehyde for 10 min before staining, image acquisition using a Leica DMI 6000 microscope, determination of the integral of the Alexa Fluor 647 intensity using FIJI^43^ and division by the integral of the intensity of nuclear mask blue for normalization. For quantifications, three randomly chosen images per well were analyzed.

### Generation of transgenic mice, immunohistochemical analysis and skeletal stainings

Mouse housing and experiments were in accordance with animal welfare laws and all relevant ethical regulations. They were approved by the responsible local committees and government bodies (Veterinäramt Stadt Erlangen, Regierung von Unterfranken). Mice carried floxed alleles for *Ep400* ^26^ or *Kat5* ^44^ in combination with a *Wnt1::Cre* transgene ^45^ for neural crest-specific deletion and a *Rosa26-stopflox-YFP* allele ^46^ for detection of the Cre-dependent recombination event. Genotyping was performed as described before ^26, 44–46^. Mice were on a mixed C3H x C57Bl/6J background and kept with continuous access to food and water under standard housing conditions in 12:12 h light–dark cycles. For bromodeoxyuridine (BrdU) labeling, 100 µg BrdU (Sigma, #B5002) per gram body weight were injected intraperitoneally into pregnant mice 1 h before tissue preparation. Embryos were obtained at E9.5, E10.5 and E18.5. For immunohistochemistry, embryos were fixed in 4% paraformaldehyde for 3-4 h (E9.5 and E10.5) or 16 h (E18.5), washed with PBS, incubated for 7 days at 4 °C in PBS + 30% sucrose, embedded in tissue freezing medium (Leica) and stored at -80 °C until further use. Sections of 10 µm thickness were produced on a Leica cryotome C1900, placed on glass slides, washed, permeabilized with 0.5% Triton X-100 and incubated in blocking solution consisting of PBS with 10% fetal calf serum, 1% bovine serum albumin and 0.5% Triton X-100 for 2 h at RT. Afterwards, sections were incubated with primary antibodies in blocking buffer at 4°C for 16 h, and then with secondary antibodies in blocking buffer for 2 h at room temperature (for antibodies see Suppl. Table 3). Sections were counterstained with DAPI and mounted with Mowiol 4-88 (Carl Roth). Image acquisition was performed with Leica DMI 6000 microscope. For staining of cartilage and bones at E18.5, embryos were fixed in 100% ethanol, incubated at room temperature for 24 h in an alcian blue solution containing 20% acetic acid, and 80% ethanol, fixed in ethanol, and consecutively stained for 48 h in an alizarin red solution containing 1% KOH. Clearing was performed in 1% KOH.

### Statistical Analysis

Results from independent animals, experiments, or separately generated samples were treated as biological replicates. Sample size was n ≥ 3 as common for this kind of study. Randomization was not possible. Investigators were not blinded in animal experiments. GraphPad Prism8 (GraphPad software, La Jolla, CA, USA) was used to determine statistical significance by multiple t-test (step-up method of Benjamini, Krieger and Yekutieli), unpaired t-test, one-/two-way ANOVA with Dunett’s post test or one-way ANOVA with Sidak’s post test as indicated in the respective figure legends (*, P ≤0.05; **, P ≤0.01; ***, P ≤0.001). The data met the assumptions of the chosen test. Variance between statistically compared groups was similar.

## Data availability

All data are provided in the manuscript and its supplement or have been submitted to GEO under accession number GSE215109.

## Acknowledgements

We thank Dirk Mielenz and Hans-Martin Jäck (Division of Immunology, Department of Medicine 3, FAU) for their help with Seahorse Measurements. R Fukunaga and G. Eichele are acknowledged for provision of Ep400 and Kat5 floxed mice. This work was supported by a grant from the IZKF Erlangen (E28).

## Contributions

Ma.W., Mi.W. and L.G. conceived the study. S.G.-B., T.S., F.F. and G.R. performed and interpreted the experiment with help of Ma.W. S.G.-B., T.S., Mi.W. and L.G. wrote the manuscript.

