## Supplemental Material for "The Tip60/Ep400 chromatin remodeling complex impacts basic cellular functions in cranial neural crest-derived tissue during early orofacial development"

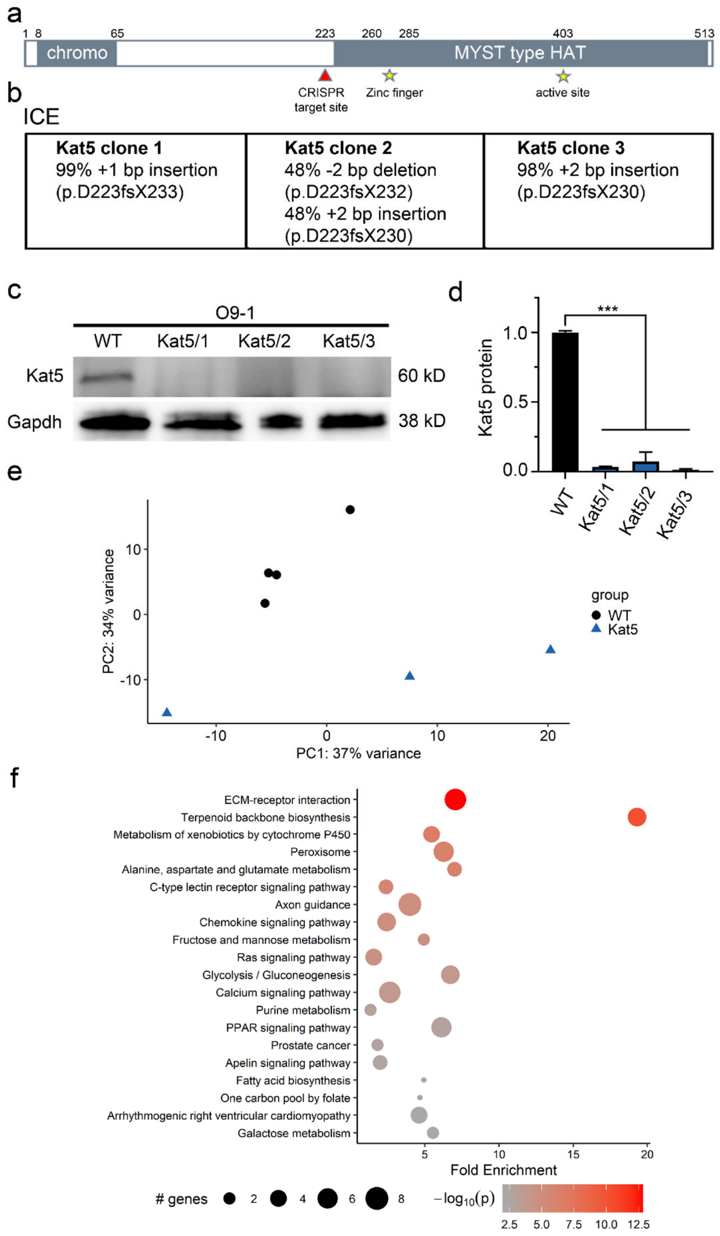


**Supplemental Figure 1: CRISPR/Cas9-dependent Kat5 inactivation in O9-1 cells. (a)** Schematic representation of the Kat5 protein, its domains (grey boxes) and structural features (asterisks) as well as the target site of the CRISPR/Cas9 editing (red arrowhead). Numbers above the bar correspond to positions of amino acid residues. Chromo, chromodomain; HAT, histone acetyltransferase domain. **(b)** Characterization of the CRISPR/Cas9-evoked gene editing events in Kat5 ko clones as determined by ICE. **(c,d)** Detection of Kat5 and Gapdh proteins in wildtype O9-1 cells (WT) and gene-edited clones by Western blotting (c) and quantification of Kat5 levels (d) relative to the amount of total protein (n=3). The size of detected proteins in kDa (left) was estimated from co-electrophoresed marker proteins. **(e)** PCA-plot of RNA sequencing samples from gene-edited clones (n=3) and wildtype cells (n=4). **(f)** KEGG clustering of genes down-regulated following Kat5 inactivation to identify gene ontology pathways and processes affected by Kat5 inactivation. Statistical significance was determined by one-way ANOVA and Dunnett's post test (***, P ≤0.001).

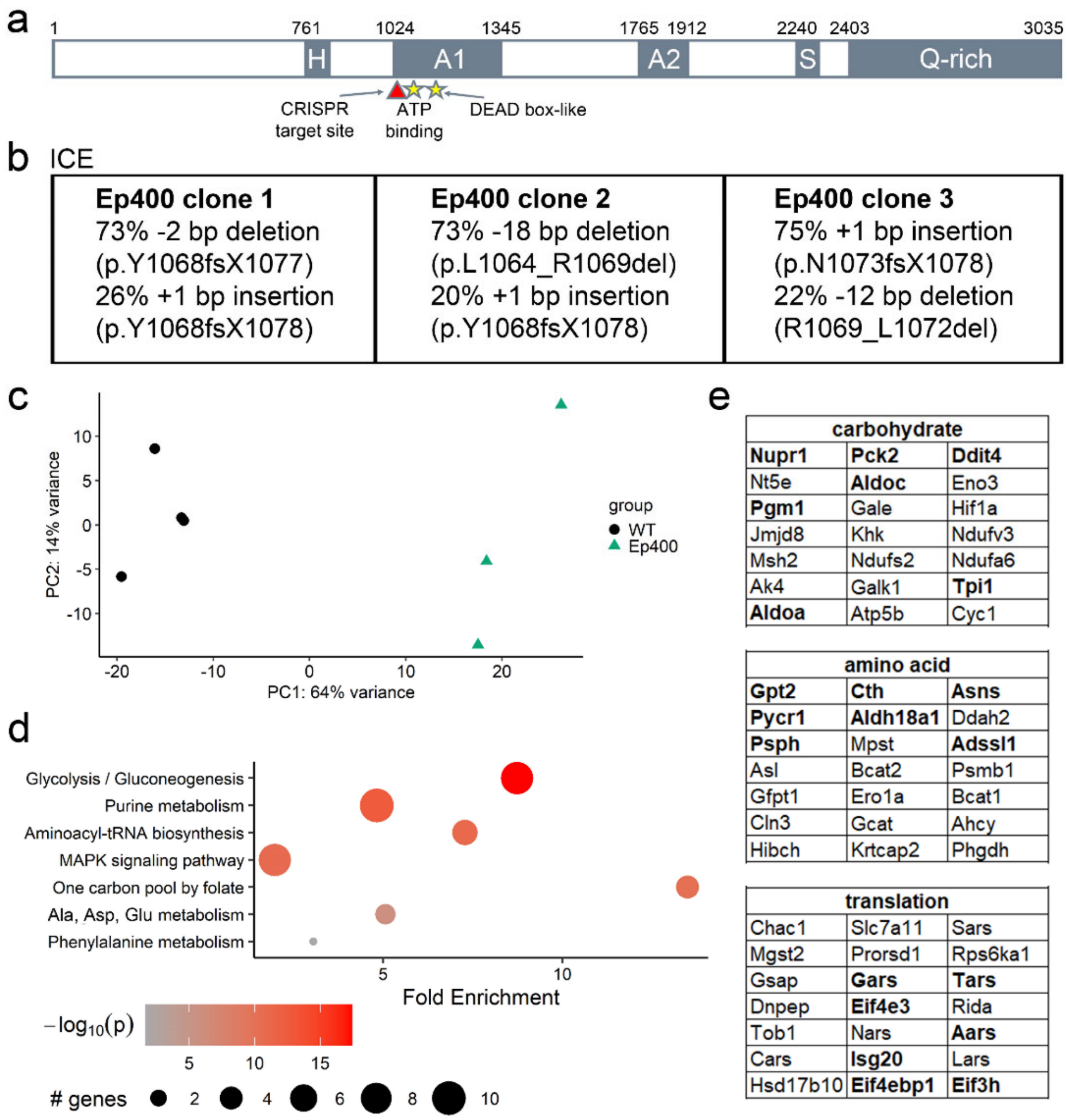


**Supplemental Figure 2: CRISPR/Cas9-dependent Ep400 inactivation in O9-1 cells. (a)** Schematic representation of the Ep400 protein, its domains (grey boxes) and structural features (asterisks) as well as the target site of the CRISPR/Cas9 editing (red arrowhead). Numbers above the bar correspond to positions of amino acid residues. H, HSA domain; A1 A2, parts of the bipartite ATPase domain; S, SANT domain; Q-rich, glutamine-rich domain. **(b)** Characterization of the CRISPR/Cas9-evoked gene editing events in Ep400 ko clones as determined by ICE. **(c)** PCA-plot of RNA sequencing samples from gene-edited clones (n=3) and wildtype cells (n=4). **(d)** KEGG clustering of genes down-regulated following Ep400 inactivation to identify gene ontology pathways and processes affected by Ep400 inactivation. **(e)** Selection of top downregulated genes shared between Ep400 and Kat5 ko clones and sorted by category (carbohydrate metabolism, amino acid metabolism, protein biosynthesis). Genes studied in validation experiments are in bold.

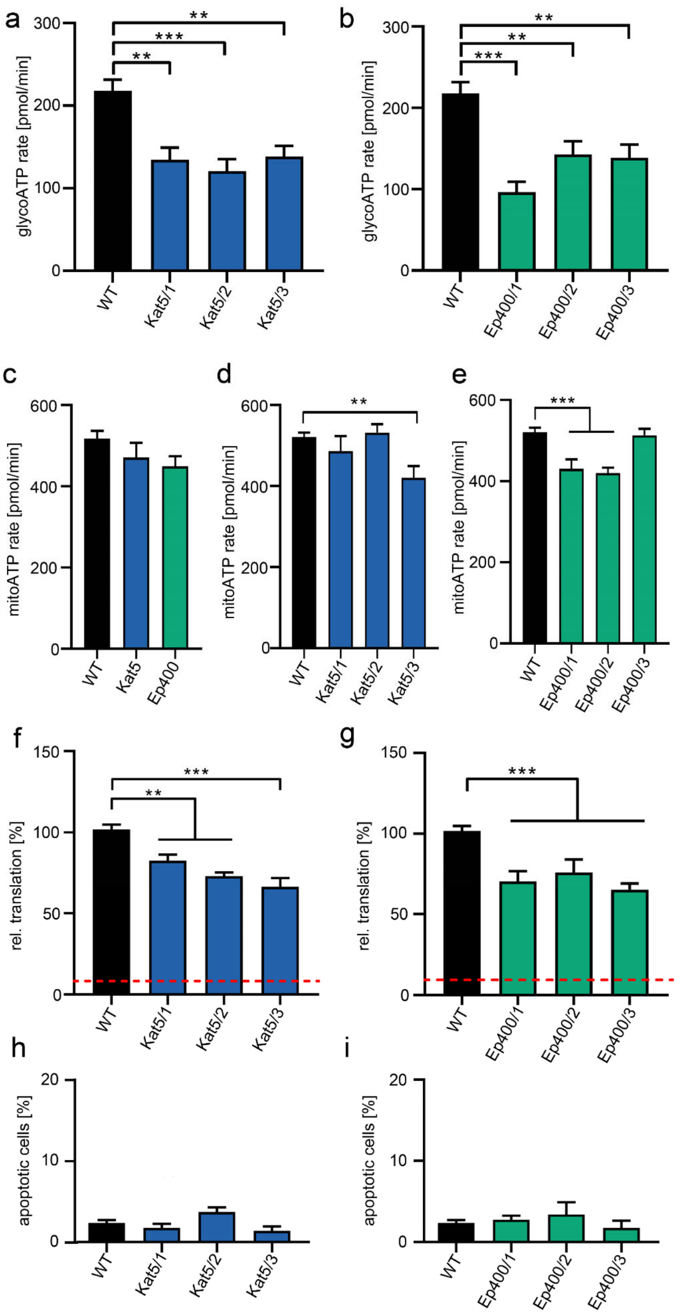


**Supplemental Figure 3: Consequences of Kat5 and Ep400 inactivation on ATP generation, translation and apoptosis in cultured O9-1 cells. (a,b)** Determination of glycolytic ATP generation rates (in pmol per minute) in single clones with Kat5 (a, blue bars) or Ep400 (b, green bars) inactivation relative to wildtype (WT, black bars) cells. **(c-e)** Determination of mitochondrial ATP generation rates (in pmol per minute) averaged over three clones with Kat5 or Ep400 inactivation (c), or determined in single clones (d, for Kat5 ko clones; e, for Ep400 ko clones) relative to wildtype cells. **(f,g)** OPP incorporation in nascent transcripts (with levels in WT cells set to 100%) in single clones with Kat5 (f) or Ep400 (g) inactivation. Red lines correspond to OPP incorporation in the presence of cycloheximide. **(h,i)** Percentage of cleaved caspase 3-positive cells in cultures of single clones with Kat5 (h) or Ep400 (i) inactivation relative to wildtype cells. Statistical significance was determined by one-way ANOVA and Dunnett's post test (*, P ≤0.05; **, P ≤0.01; ***, P ≤0.001).

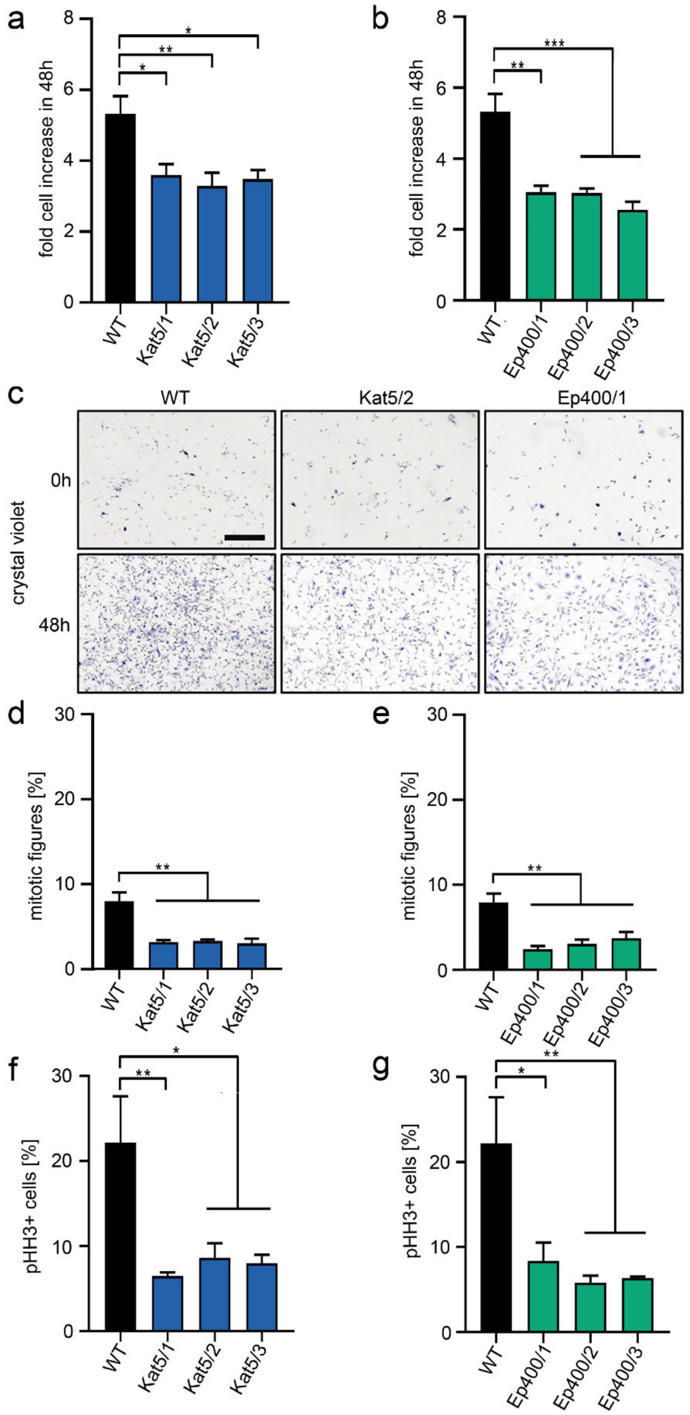


**Supplemental Figure 4: Consequences of Kat5 and Ep400 inactivation on proliferation in cultured O9-1 cells. (a,b)** Determination of 48 h proliferation rates in single clones with Kat5 (a, blue bars) or Ep400 (b, green bars) inactivation relative to wildtype (WT, black bars) cells as determined by crystal violet staining. Staining levels at 0 h were set to 1. **(c)** Representative images of crystal violet stainings of cultured wildtype, clone Kat5/2 and clone Ep400/1 cells at 0 h and 48 h. Scale bar: 400 µm. **(d-g)** Percentage of mitotic (d,e) and phosphohistone H3-positive (f,g) cells in cultures of single clones (d,f, for Kat ko clones; e,g, for Ep400 ko clones) relative to wildtype cells. Statistical significance was determined by one-way ANOVA and Dunnett's post test (*, P ≤0.05; **, P ≤0.01; ***, P ≤0.001).

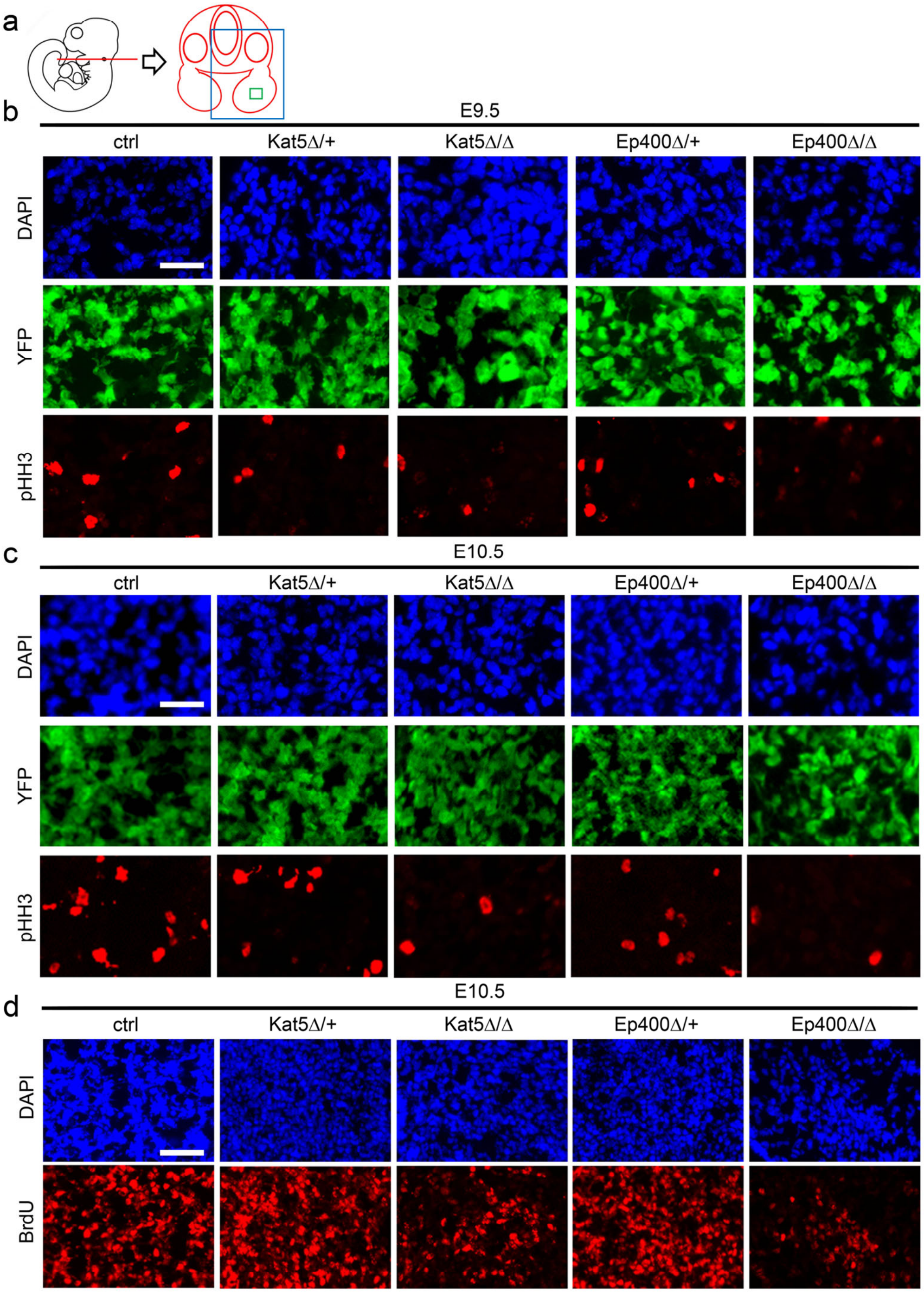


**Supplemental Figure 5: Consequences of Kat5 and Ep400 inactivation on cranial neural crest proliferation. (a)** Schematic representation of early mouse embryo and section used for analyses (red). The larger boxed area (blue) is depicted in Fig. 6c, the contained smaller area (green) in Suppl. Figs 5b-d and 6.a,b **(b-d)** Representative images of immunohistochemical stainings with anti-phosphohistone H3 (b,c, pHH3, red, bottom row) or anti-BrdU antibodies (d, red, bottom row) of transverse sections of the first pharyngeal arch from control (ctrl) embryos and age-matched heterozygous as well as homozygous embryos with Kat5 (Kat5Δ/+, Kat5Δ/Δ) or Ep400 (Ep400Δ/+, Ep400Δ/Δ) deletions at E9.5 (b) or E10.5 (c,d). DAPI counterstains are shown in the top row (blue), the neural crest-specific YFP autofluorescence in the middle row (green). Scale bar: 50 µm (b,c), 100 µm (d).

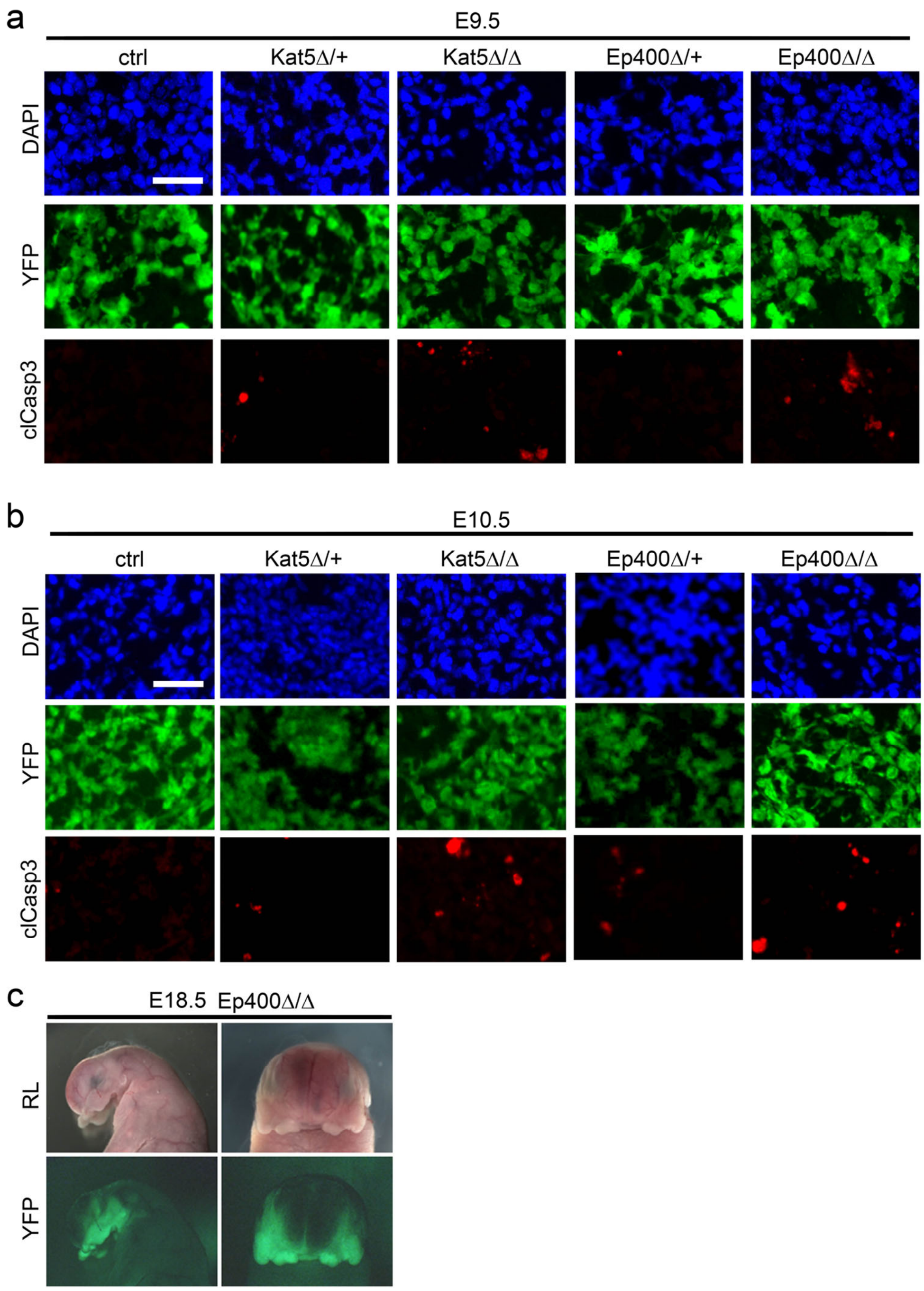


**Supplemental Figure 6: Consequences of Kat5 and Ep400 inactivation on cranial neural crest survival. (a,b)** Representative images of immunohistochemical stainings with anti-cleaved caspase 3 antibodies (red, bottom row) of transverse sections of the first pharyngeal arch from control (ctrl) embryos and age-matched heterozygous as well as homozygous embryos with Kat5 (Kat5Δ/+, Kat5Δ/Δ) or Ep400 (Ep400Δ/+, Ep400Δ/Δ) deletions at E9.5 (a) or E10.5 (b). DAPI counterstains are shown in the top row (blue), the neural crest-specific YFP autofluorescence in the middle row (green). Areas for presentation were as in Suppl. Fig. 5a. Scale bar: 50 µm. **(c)** Reflected-light microscopic (RL, upper row) and YFP-autofluorescent (lower row) images of heads from homozygous embryos with Ep400 (Ep400Δ/Δ) deletions at E18.5 (for comparison to ctrl and Kat5Δ/Δ embryos, see Fig. 7a.
